# The *Drosophila* Neurogenin, Tap, controls axonal growth through the Wnt adaptor protein Dishevelled

**DOI:** 10.1101/034439

**Authors:** Liqun Yuan, Shu Hu, Zeynep Okray, Xi Ren, Natalie De Geest, Annelies Claeys, Jiekun Yan, Eric Bellefroid, Bassem A. Hassan, Xiao-Jiang Quan

## Abstract

The Neurogenin (Ngn) transcription factors control early neurogenesis and neurite outgrowth in mammalian cortex. In contrast to their proneural activity, their function in neurite growth is poorly understood. *Drosophila* has a single predicted Ngn homologue called Tap, whose function is completely unknown. Here we show that Tap is not a proneural protein in *Drosophila* but is required for proper axonal growth and guidance of neurons of the mushroom body (MB), a neuropile required for associative learning and memory. Genetic and expression analyses suggest that Tap inhibits excessive axonal growth by fine regulation of the levels of the Wnt signaling adaptor protein, Dishevelled.

**Summary:** The *Drosophila* Neurogenin homologue, Target of Pox neuro (Tap), prevents axonal overgrowth by regulating the Wnt Planar Cell Polarity pathway adaptor protein Dishevelled.

## Introduction

Proper function of the nervous system is based on the production of a diversity of neuronal and glial cells, as well as the precise targeting of their axons and dendrites. A key transcription factor (TF) family, which controls the commitment of neuronal cell fate and neurite guidance, is the Neurogenin (Ngn) family. Ngn proteins belong to a structurally and functionally conserved basic helix-loop-helix (bHLH) superfamily. Ngns in vertebrates are sufficient to initiate the neuronal cell fate in the central nervous system (CNS) (reviewed in Bertrand et al., 2002; Ma et al., 1998). In contrast, very little is known about the function of Ngn proteins in invertebrate systems. Gain of function of analyses in *Drosophila* and vertebrate models suggested that during evolution a switch in proneural activity occurred between the Ngns and a highly related family of bHLH TFs called the Atonal family. Specifically, whereas Ngns are necessary and sufficient for the induction of neurogenesis in vertebrates, they cannot do so in flies. Conversely, in flies Atonal type proteins can induce neurogenesis, but fail to do so in vertebrates (Quan et al., 2004). A crucial test of this “evolutionary proneural switch hypothesis” is whether or not *Drosophila* Neurogenins act as proneural genes in flies and vertebrates.

In addition to their proneural function, Ngns play various critical roles in the development of vertebrate nervous system, including regulating the outgrowth and targeting of both axons and dendrites (Hand et al., 2005; Hand and Polleux, 2011; reviewed in Yuan and Hassan, 2014). The regulation of such neurite growth is independent from the proneural activity, as the phosphorylation of a single tyrosine in mouse Ngn2 is necessary to specify the dendritic morphology without interfering the cell fate commitment (Hand et al., 2005). However, the mechanism of how Ngn proteins regulate neurite guidance is poorly understood.

The *Drosophila* genome encodes a single member of Ngn family based on the conservation of family-defining residues in the bHLH domains (Hassan and Bellen, 2000), named Target of Pox neuro (Tap). Previous work suggests that Tap is expressed in secondary progenitors of both neurons and glia, called ganglion mother cells, in CNS (Bush et al., 1996), as well as the putative support cells in the peripheral nervous system (PNS) during embryogenesis (Gautier et al., 1997). A previously reported putative *tap* mutant allele (Ledent et al., 1998) was later shown to be a mutation in a different gene called *blot* (Johnson et al., 1999). Therefore, the function of the *Drosophila* Ngn Tap remains uncharacterized.

Here we generated a null mutant allele of *tap* by replacing the single coding exon of the *tap* gene with Gal4. Using expression, gain of function and loss of function analyses we show that, whereas ectopic expression of Tap in vertebrates can induce neurogenesis, *tap* is not a proneural gene in flies, consistent with the evolutionary proneural switch hypothesis. Instead, Tap is required to prevent overgrowth of axons during brain development, at least in part through the activity of the axonal Wnt planar cell polarity (Wnt-PCP) pathway by fine tuning the levels of the Wnt signalling adaptor protein Dishevelled (Dsh).

## Results and Discussion

### Tap is the only Ngn homolog in *Drosophila*

*Drosophila* Tap shares significant identity (~70%) in the bHLH domain with mouse and human Ngns (**Fig. 1A**), compared to *Drosophila* Atonal or Scute. To test if Tap, like its vertebrate courter parts, has proneural activity, we injected *Tap* mRNA into one cell of 2-cell stage *Xenopus* embryos and assessed neurogenesis by staining for neuronal markers. Like mouse Ngn1, but unlike fly Atonal, Tap efficiently induces neurogenesis in this system (**Fig. 1B-E**). Conversely, when ectopically expressed in a classic *Drosophila* proneural assay Tap – like Ngn1, and in contrast to Atonal – fails to induce neurogenesis (**Fig. 1H-I**). These data suggest that Tap may not be a proneural gene in flies, even though it has neural induction potential in vertebrates, as expect for a bona fide Ngn protein.

**Figure 1.**
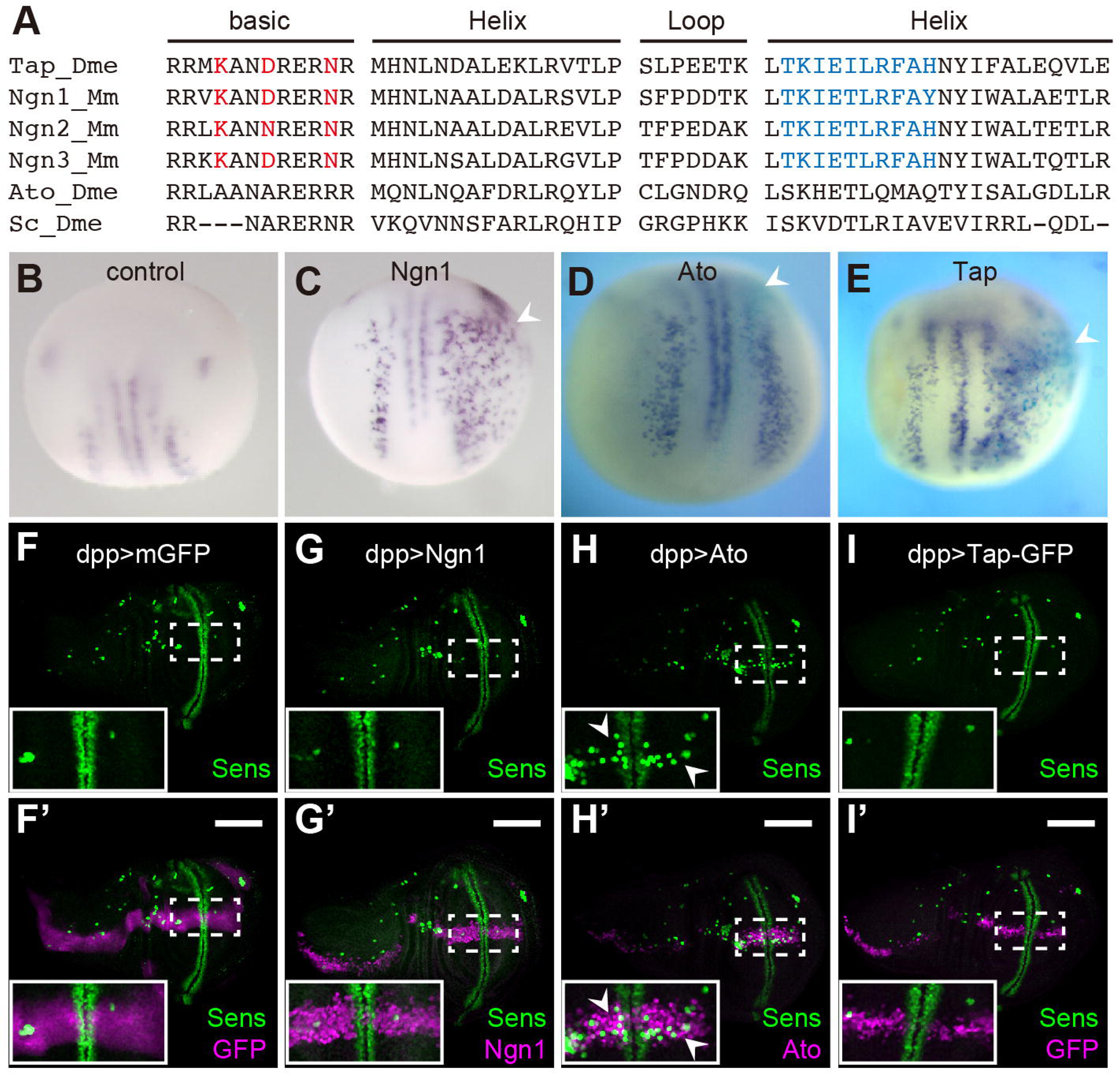
Tap is the Ngn homolog in *Drosophila*. **A**. Sequence comparison among bHLH domain of fly Tap, mouse Ngn family, fly Ato and Scute. The three residues in basic domain (red) were identified as defining the proneural function of Ngn and Ato in vertebrates and flies respectively. The ten residues in the second Helix domain (blue) are also class-specific (Quan et al., 2004). Dme: *Drosophila Melanogaster*; Mm: *Mus musculus*. **B**-**E**. *In situ* hybridization of N-tubulin is used to visualize neurogenesis in *Xenopus* embryos upon no injection (**B**), mRNA of mouse Ngn1 (**C**), Ato (**D**) or Tap (**E**), (arrowhead). **F**-**I**’. The distribution of neuronal precursors (Sens, green) in L3 wing discs. CD8::GFP (**F**), mouse Ngn1 (**G**), Ato (**H**) or Tap-GFP (**I**) is ectopically expressed in the anterior-posterior axis of wing discs (magenta), driven by *dpp-Gal4*. Scale bars=100 μm.

### Tap is widely expressed in the nervous system during development

To investigate the function of Tap, we generated a mutant allele using ends-in homologous recombination (Rong and Golic, 2000) to replace the open reading form of *tap* with an external driver, Gal4 (**Fig. 2A**). The *tap ^Gal4^* allele, in homozygosity, serves as a *tap* null mutant; while in heterozygosity, it was used as a driver to reveal the expression pattern of Tap. We made several attempts to generate a Tap antibody, but this was not successful. Tap^Gal4^ was detected in a large number of cells in both the central and peripheral nervous systems throughout development (**Fig. 2B-E**). During embryogenesis, Tap is enriched in the ventral nerve cord (VNC) and sparsely expressed in the PNS (**Fig. 2B, S2**). Postembryonically, Tap positive cells were observed in many tissues, including optic lobes (OLs), MBs, antenna lobes (ALs) and subesophageal ganglion (SOG) (**Fig. 2C-E**).

**Figure 2.**
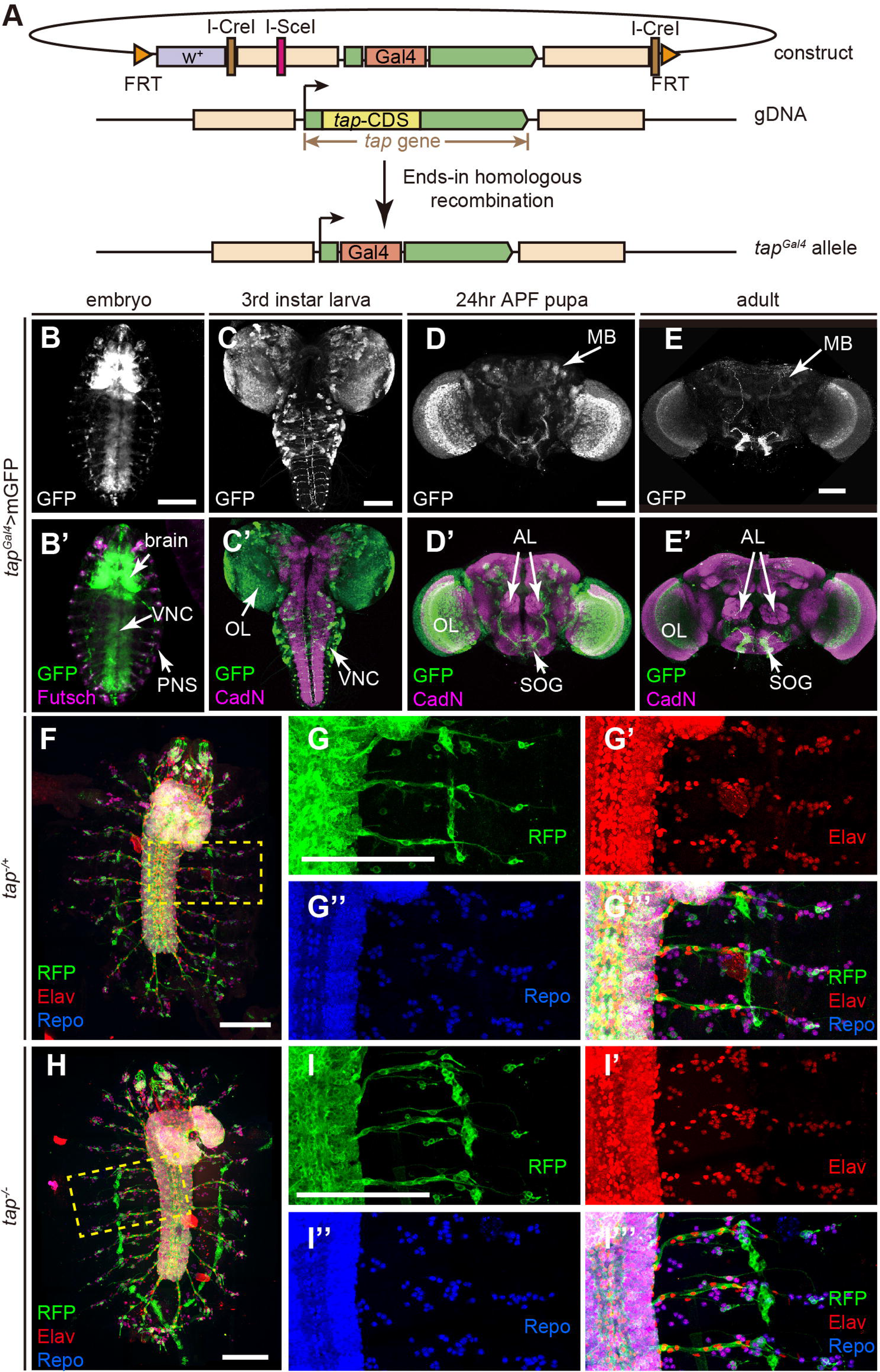
Characterization of Tap expression pattern. **A**. Schematic diagram of generation of *tap^Gal4^* allele. **B-E**. Tap is widely expressed in nervous system throughout development. Tap^+^ cells were labelled by membrane GFP (mGFP) driven by *tap^Gal4^*. **B**. Ventral view of a stage 16 embryo. VNC: ventral nerve cord. **C**. A L3 brain. OL: optic lobe. **D**. Brain of a 24h APF pupa. MB: mushroom body; AL: antenna lobe; SOG: subesophageal ganglion. **E**. An adult brain. **F-I**. Flat preparation of whole embryos at stage 17. Tap^+^ cells are labelled with CD8::RFP. The differentiated nervous system is visualized as neurons (red) and glia cells (blue). **F**. *tap* heterozygous (*tap^Gal4^/+*) embryo. **G**. Magnification of the region framed in **F**. **H**. *tap* null (t*ap^Gal4/Gal4^*) embryo. **I**. Magnification of the framed region in **H**. Scale bar=100 μm.

### Tap is not required for specifying the neuronal or glial cell fate during embryogenesis

Flies lacking Tap are mostly embryonic lethal, with a few escapers to early larval stages. This lethality can be rescued, including to adult viability, by re-expression of Tap (*UAS::tap*) in *tap^Gal4^* homozygous flies. We examined whether the number or fate of neuronal and/or glial cells are altered in *tap* mutant embryos. Surprisingly, we find no obvious morphological defects in the *tap* mutant embryos. The number and pattern of the neurons and glia in the PNS are intact. Although the cell number in the CNS is difficult to quantify, the general cellular pattern looks similar in the mutant when compared with the control flies (**Fig. 2F-I**). These data suggest that Tap is not required for early neurogenesis in *Drosophila* embryos.

***tap* mutants show MB β lobe midline-crossing and α lobe missing defect**

To characterize Tap function, we focused on the MB as a model system. Tap is expressed in the MB at both pupal and adult stages in a subset of α/β neurons (**Fig. 3A-C**), which form four clusters in the cell body (**Fig 3C**) and project their medial axons to the dorsal part of β lobe (**Fig 3A, B**). Tap expression is highest at early pupal stages, and then declines. The phase of high levels of Tap expression correlates with the differentiation stage of α/β neurons, indicating Tap may regulate the development of the α/β lobe.

**Figure 3.**
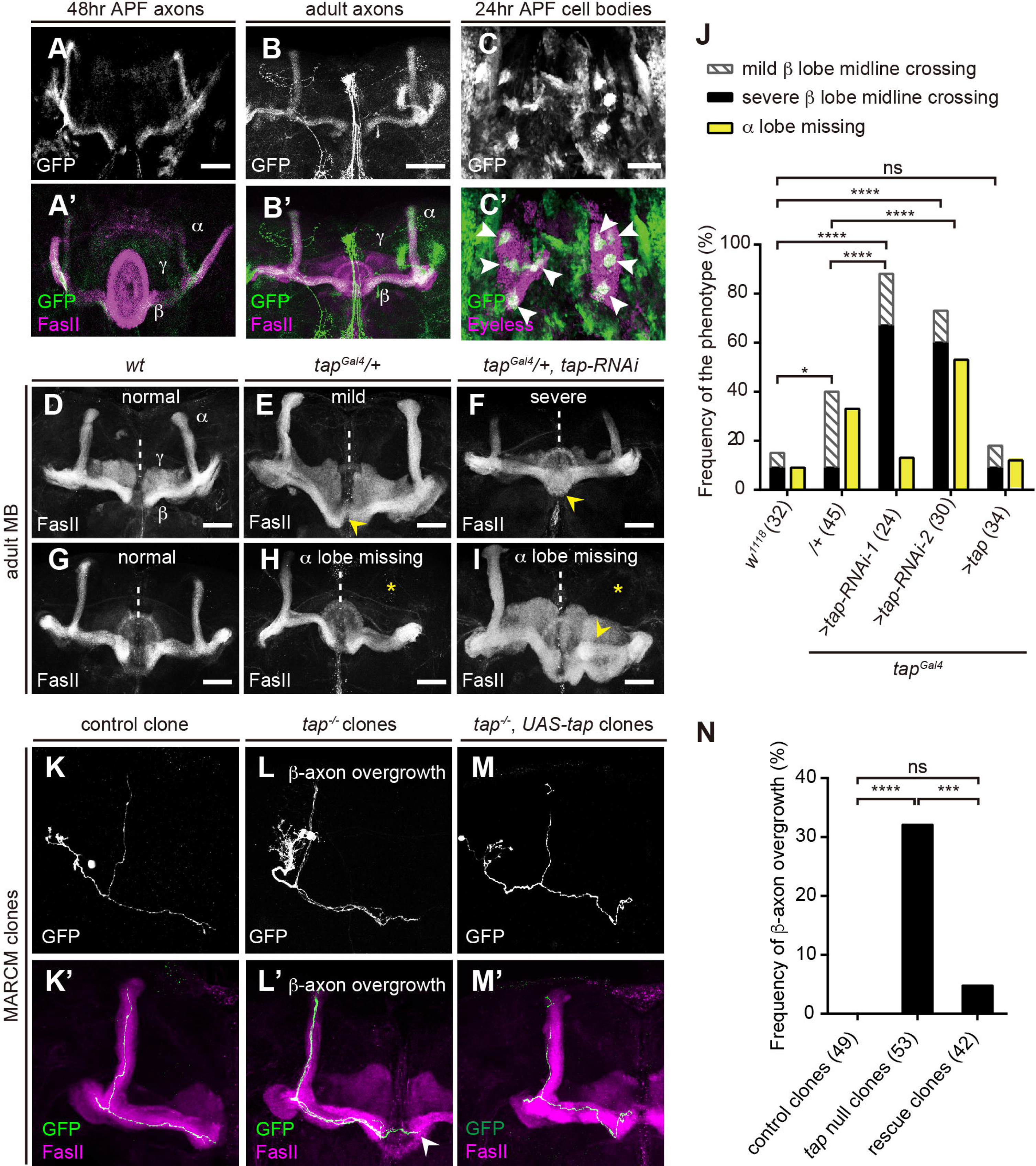
Tap is required for the axonal growth and guidance of MB α/β neurons. **A**-**B**. Z projection of MB axon level at 48h APF pupa (**A**) or adult (**B**). FasII (magenta) strongly labels the α and β lobes, and weakly labels γ lobes (**A’**, **B’**). GFP expression is restricted to a subset of axons in the distal region of α lobes and ventral region of β lobes. **C**. GFP is concentrated in four clusters of cells (arrowhead) within MB cell bodies (magenta). **D**. Normal MBs in a wild-type fly. **E**. A *tap* heterozygous mutant brain exhibits mild β axon midline crossing defect in which the thickness of the fiber bundle crossing the midline is less than the width of the β lobe termini (arrowhead). **F**. A tap knock-down MB using RNAi appears severe β lobe fusion defect in which the crossing fibers equal in width to the terminal of adjacent β lobes (arrowhead). **G**. Normal MBs in a wild-type fly. **H-I**. *tap* mutants lose the dorsal projection of α/β neurons (asterisks). The loss of dorsal lobe is sometimes accompanied by the bifurcation of the medial lobe (arrowhead in **I)**. **J**. Fraction of brains that displays the defects. Numbers of the brains being analysed were indicated in the bracket. ns: p=0.9242; *: p=0.0280; ****: p<0.0001, Chi-squared test. **K**-**M**. One half of the adult MBs. Axonal projections of labelled MB neurons generated by MARCM were visualized by GFP driven by *ey^OK107^-Gal4*. **K**. A control clone. **L**. *tap* null clones. β axons exceed the region of β lobe, extending to the contralateral β lobe region (arrowhead). **M**. A rescue clone in which Tap is re-introduced in the mutant clonal cell only. **N**. Fraction of clones that displays the defect of β-axon overgrowth. Numbers of the clones being analysed were indicated in the bracket. ns: p=0.1224, ****: p<0.0001, ***: p=0.0009, Chi-squared test. Scale bar =50 μm.

To circumvent embryonic lethality we began by exploiting the heterozygous *tap ^Gal4^* allele alone or in combination with two independent RNAi strains targeting distinct regions of *tap*. In wild-type *Drosophila*, axons of the medially projecting β lobes terminate near the midline but do not cross it (Strausfeld et al., 2003). In contrast to the wild-type MB morphology (**Fig. 3D**), *tap* loss-of-function brains exhibit that β lobe fibers extend across the midline (**Fig. 3E**), sometimes sufficient to cause fusion of the two contralateral β lobes (**Fig. 3F**). The phenotype is variable in severity, thus we classified it as “normal”, “mild”, or “severe” based on the thickness and density of the β lobe fibers crossing the midline. Tap heterozygotes display an increase of the penetrance of mild defects, while both *tap* RNAi strains raise the incidence and severity of lobe midline crossing defect. When Tap is re-expressed, the defect can be rescued to control levels (**Fig. 3J**). This β axon midline-crossing defect is developmental in origin as it can be observed at early pupal stage (**Fig. S3**).

In addition to the β axon midline-crossing defect, an “α lobe missing” phenotype was observed in Tap loss of function MBs (**Fig. 3H, I**). In some brains with a missing α lobe, the β lobe appears to branch into two bundles (**Fig. 3I**), suggesting this α lobe defect may be an axonal targeting defect rather than a growth defect. Unlike the β lobe defect, the penetrance of α lobe missing phenotype varies considerably (**Fig. 3J**). Considering the frequency is still higher in the *tap* mutant flies compared to wild-type and rescue strains, we conclude that Tap is essential for the growth and correct targeting of both lobes of α/β neurons.

### Tap is required cell-autonomously for the growth and targeting of the β lobe

Since the branching and growth pattern of single α/β axons cannot be directly inferred from the morphology of the α/β lobes, a single-neuron level analysis, such as Mosaic analysis with a repressible cell marker (MARCM), is necessary to clarify the targeting of α/β neurons (Lee et al., 2000). Therefore *tap* null clones were generated in a *tap^Gal4^* heterozygous background using the MARCM technique. In both wild-type control and *tap^Gal4^* heterozygous background, a small minority of brains showed β lobe overgrowth and/ or α lobe missing defect as discussed above. Considering this, only brains with intact over all α/β lobes morphology were quantified for clonal axon phenotypes. Analysis of small MB clones reveals that none of the control clones show any defects (**Fig. 3K**). In contrast, in 32% of *tap* null mutant clones, β axons project beyond the β lobe domains (**Fig. 3L, N**). In severe cases, axons were observed to cross the midline and project to the contralateral β lobes. This defect can be rescued by re-introduction of *tap* specifically in the mutant clones (**Fig. 3M, N**). This suggests Tap is required cell-autonomously for the development of the β axon branch. However, none of the mutant clones showed loss of α axon growth, suggesting Tap plays a non-cell-autonomous role during α lobe development.

### Tap regulates axon growth through Dsh

In order to investigate the molecular mechanism of how Tap regulates axonal growth, we performed dominant interaction tests between Tap and well-established MB axonal guidance factors, particularly those whose loss of function causes β lobe overgrowth. Specifically, we asked if heterozygosity for any of these genes strongly enhances the very mild phenotypes observed in *tap* heterozygotes over the sum of the phenotypes observed in each allele. Among the candidate genes, *drl*, *Sli* and *Dscam1* did not show any obvious synergism with Tap (**Fig. 4A**). However, loss of one copy of *dsh* strongly enhances both α lobe guidance and β lobe overgrowth defects in the *tap* heterozygous background (**Fig. 4A, B**). This suggests that Dsh synergizes with Tap to regulate the axon guidance and growth during MB development. To test if Tap might regulate Dsh expression *in vivo* we overexpressed Tap or knocked it down in all neurons and measured Dsh protein levels. We find that Tap gain of function mildly increased Dsh levels, while *Tap* knockdown mildly decreases Dsh levels *in vivo* (**Fig. 4C, D**), although these changes are not strong enough to reach statistical significance.

**Figure 4.**
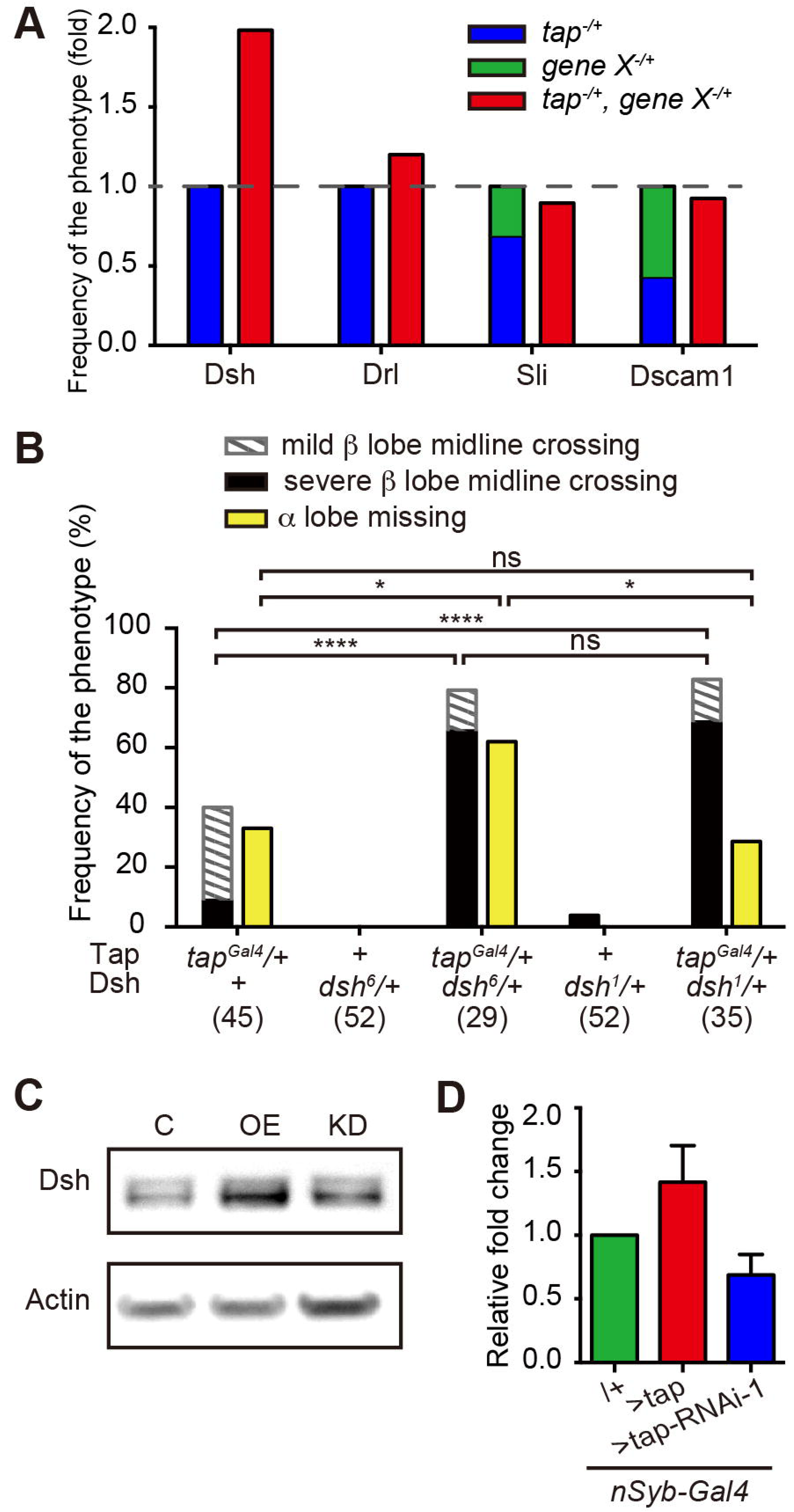
Tap regulates axon guidance through Dsh in PCP pathway. **A**. Genetic interactions for the defect of β-lobe overgrowth in double heterozygous mutant between Tap and axon guidance factors known to cause β-lobe overgrowth. The penetrance of total β lobe overgrowth defect was normalized to the sum of the penetrance of *tap* mutant plus the mutant of candidate genes. Dashed line indicates 1, which represents the normalized defect severity aggregated by the two single mutants. **B**. Fraction of brains that displays the defects in *tap* and *dsh* double heterozygous mutant. Numbers of the brains being analysed were indicated in the bracket. *dsh^1^* is a PCP pathway specific mutant of *dsh*, and *dsh^6^* is the loss-of-function mutant of *dsh*. **C**. Western blot analysis showing Dsh level in driver alone (C), Tap overexpression (OE) or knockdown (KD) cases. Actin is used as a loading control. **D**. Quantification of relative changes in Dsh level of OE or KD, normalized by control. Error bars represent the s.e.m., n=3.

Dsh is a key component in Wnt signaling pathway, which serves as a hub to relay the signals from the receptors to the downstream effectors (Gao and Chen, 2010). It is essential in both canonical and non-canonical Wnt pathway, such as Wnt-PCP pathway. In *Drosophila*, PCP pathway is known to regulate distinct phenomena including polarity within cell sheets (Fanto and McNeill, 2004), dendritic arborization (Gao et al., 2000) and axonal growth and guidance (Ng, 2012) and the role of Dsh in MB β lobe axonal growth is mediated by its activity in the Wnt-PCP pathway (Shimizu et al., 2011). To further query the interaction between Tap and Dsh we took advantage of the Wnt-PCP specific allele of Dsh in Drosophila, *dsh^1^*. Interestingly, the combination of *dsh^1^* and *tap* heterozygousity increases the severity of β lobe overgrowth defect to a similar level as double heterozygote of *tap* and the null allele of *dsh*, *dsh^6^*, but does not affect the α lobe missing defect (**Fig. 4B**). These data suggest that the role of Tap in the regulation of axonal growth and guidance in α lobe and β lobe are independent. Furthermore, this is consistent with previous findings that Wnt-PCP signaling β lobe growth cell autonomously, but α lobe growth cell non-autonomously (Soldano et al., 2013). In addition to Dsh, three other key Wnt-PCP components regulate αβ lobe growth namely, Appl and Vang and Wnt5a. We find that heterozygosity for all three mildly enhances the *tap* defect, but much less than Dsh does, suggesting that Tap regulates PCP signaling specifically through Dsh (**Fig. S4**).

Our data suggest that *tap* regulates neuronal extension and guidance through the Wnt-PCP pathway. Cellular polarity created by the PCP pathway is essential to direct cell movement, which is analogous to guide the movement of growth cones, thus navigate the axonal growth (Zou, 2012). The regulation of PCP pathway was characterized via protein-protein interactions previously. For instance, Vangl2, one core receptor of PCP pathway in mammalian, is found to antagonize with Disheveled, the mammalian homolog of *Drosophila* Dsh, and posttranslationally regulate another receptor protein, Frizzled3, to transmit the information to the downstream target genes of PCP signaling pathway, in such a way to regulate the polarity of growth cones in the filopodia (Shafer et al., 2011). Here we identify a transcriptional modulator of the PCP pathway during axonal growth, although it is unclear whether Dsh is a direct or indirect target of Tap. The changes in Dsh protein levels upon Tap manipulation are quite mild suggesting that Tap does not serve as an activator, but as an enhancer of the PCP pathway. This, however, is consistent with the observation that double heterozygosity for *tap* and *dsh* phenocopies the strongest knock down of *tap*. This implies that the neuronal Wnt-PCP pathway is sensitive to the quantitative regulation of Dsh by Tap.

In contrast to the potential conservation of its function in neurite growth, Tap does not act as proneural gene in flies. Despite of the obvious differences in morphology between invertebrates and vertebrates, the fundamental mechanisms underlying neurogenesis are conserved. In particular, the induction of neurogenesis by bHLH proteins has been conceptually defined as a module that can be applied to various contexts (Schlosser and Wagner, 2004). Our data, together with previous studies, suggest that during the split between vertebrates and invertebrates, the neurogenic module exchanged one of its major components resulting in a switch in the sensitivity of the vertebrate, but invertebrate neuroectoderm to the neuroinductive activity of Neurogenin proteins. Crucially, this is not encoded as a change in the inductive capacity of the invertebrate Neurogenins per se, as demonstrated by Tap’s neurogenic activity in *Xenopus*. Whether this is related to the dorso-ventral axis switch in the location of the neuroectoderm (dorsal in vertebrates, ventral in *Drosophila*) would be interesting to determine.

## Material and Methods

A brief description of the material and methods is presented here. Please see supplemental for detailed experimental procedures.

### Fly Husbandry and Transgenic Lines

Flies were kept at 25°C or 29°C on standard medium. Experiments involving RNAi technique, or genetic interaction were performed at 29°C, and the others were performed at 25°C.

### *Xenopus* embryo microinjection

The mRNAs of *ato*, *tap* and mouse *Ngn1* were injected in a single blastomere of *Xenopus* embryos at the two-cell stage. The whole embryos were *in situ* hybridized with anti-N-tubulin as described in (Quan et al., 2004).

### Cloning and gene targeting

The fragments of 5’ homologous recombination arm of *tap* ORF, Gal4 and 3’ recombination arm of *tap* ORF were amplified and subcloned into pED13(M) vector (gifted from B. Dickson lab). *tap* targeting was achieved by ends-in homologous recombination described in Fig. S1 and (Rong and Golic, 2000).

### Immuohistochemistry

Embryos and dissected tissues were stained using the protocol described in (Hassan et al., 2000; Langen et al., 2013). Flat preparation of embryonic fillets at stage 17 were generated and stained on polylysine-coated glass following the protocol described in (Featherstone et al., 2009). Pictures were acquired using LEICA TCS SP8 or LEICA TCS SP5 and processed with ImageJ.

### Western blot

Proteins from 20 adult fly brains in each genotype were resolved and probed using the protocol described in (Okray et al., 2015). Primary antibodies were used as: anti-Dsh 1:1000 (gifted from C. Bagni lab) and anti-Actin 1:5000 (Abcam).

## Acknowledgements and Funding

LY acknowledge discussion with A. Soldano and M. Nicolas, as well as G. Linneweber for the technical help of flat preparation of embryo samples. This work was supported by Fonds Wetenschappelijke Onderzoeks [G050312 to X.Q and B.A.H.], by the Vlaams Instituut voor Biotechnologie [to B.A.H.], by a doctoral fellowship from China Scholarship Council [2010638041 to L.Y.] and a grant from the National Natural Science Foundation of China [31171256 to S.H.]. XR was a research fellow from the FRIA.

## Author contributions

LY conceived and designed the study, collected and analyzed the data, wrote the manuscript. SH generated the mutant of *tap^Gal4^* and collected data. ZO helped with the genetic interaction screen. XR performed the microinjection of *Xenopus* embryos. NDG carried out the western blot analyses. AK and JY carried out the cloning and provided technical assistance. EB conceived and guided the gain of function experiments. BAH and XQ conceived and supervised the study, and wrote the manuscript. All authors read and approved the final manuscript.

